# Musical expertise is associated with improved neural statistical learning

**DOI:** 10.1101/2020.05.20.106187

**Authors:** Jacques Pesnot Lerousseau, Daniele Schön

**Author notes:** **Corresponding Author and Lead Contact:** Jacques Pesnot Lerousseau, Aix-Marseille Univ, INS, Inst Neurosci Syst, Marseille, France. **Author contributions:** Conceptualization J.P.L. and D.S.; Data curation J.P.L.; Formal Analysis J.P.L.; Funding acquisition D.S.; Investigation J.P.L.; Methodology J.P.L. and D.S.; Project administration D.S.; Resources D.S.; Supervision D.S.; Visualization J.P.L.; Writing – original draft J.P.L. and D.S.; Writing – review & editing J.P.L. and D.S.

## Abstract

It is poorly known whether musical training leads to improvements in general cognitive abilities, such as statistical learning (SL). In standard SL paradigms, musicians have better performances than non-musicians. However, these better performances could be due to an improved ability to process sensory information, as opposed to an improved ability to learn sequence statistics. Unfortunately, these very different explanations make similar predictions on the performances averaged over multiple trials. To solve this controversy, we developed a Bayesian model and recorded electroencephalography (EEG) to study trial-by-trial responses. Our results confirm that musicians perform ~15% better than non-musicians at predicting items in auditory sequences that embed either simple or complex statistics. This higher performance is explained in the Bayesian model by parameters governing SL, as opposed to parameters governing sensory information processing. EEG recordings reveal a neural underpinning of the musician’s advantage: the P300 amplitude correlates with the Bayesian model surprise elicited by each item, and so, more strongly for musicians than non-musicians. Finally, early EEG components correlate with the Bayesian model surprise elicited by simple statistics, as opposed to late EEG components that correlate with Bayesian model surprise elicited by complex statistics surprise, and so more strongly for musicians than non-musicians. Overall, our results prove that musical expertise is associated with improved neural SL, and support music-based intervention to fine tune general cognitive abilities.

## Introduction

Musical training provides a rich temporal structure, which is thought to improve the ability to detect and use temporal regularities. Although temporal regularities can be represented at different levels of details ^1^, the simplest representation of the sequence is the knowledge about its summary statistics. The ability to learn sequence statistics is referred to as “statistical learning” (SL). Having improved SL abilities would allow musicians to make more accurate inferences about future events, thus improving perception ^2^, decision-making ^3^, and language acquisition ^4–6^. In this definition, statistics is employed in a broad sense, and can refer for example to the frequency of occurrence of individual items (*e.g.* the frequency of ⚫, P(⚫), in ⚫⚫⚪⚫⚫⚫⚪) or to the transition probability between items (*e.g.* the transition probability of ⚫➞⚪, P(⚪|⚫), in ⚫⚪⚪⚫⚪⚫⚪). In reference to discrete Markov chain analysis, those statistics reflect different orders of Markov chains. The probability of occurrence of an item given the preceding one, *e.g.* P(⚪|⚫), is known as 1^st^ order Markov probability. Similarly, the probability of an item given the preceding K items, *e.g.* P(⚪|⚫⚪), is known as K^th^ order Markov probability. The concept of Markov chain order defines an ordering of SL, from simple to complex as K increases.

Musicians have better performances than non-musicians in tasks that implicate different levels of SL complexity. For example, musicians have a greater neural sensitivity to the frequency of occurrence of items than non-musicians ^7^, they are better at segmenting words from an artificial language stream ^8^, they have a greater neural sensitivity to 1^st^ order Markov probability ^9^ and to more complex statistics ^10–12^. However, it is not clear whether these differences arise from an improved ability to learn sequence statistics — and if so, at which level of complexity — or from an improved ability to process sensory information. Unfortunately, these alternative explanations make similar predictions on the performances averaged over multiple trials. Furthermore, computational models used to study trial-by-trial responses ^13–16^ were developed to uncover the general principles of SL. Being general by nature, they do not incorporate individual parameters, and are thus unable to account for inter-individual differences ^17,18^. To solve these problems, we created a Bayesian model and recorded electroencephalography (EEG) to both study trial-by-trial responses and account for inter-individual differences.

We asked the following questions: do musicians have better abilities to predict items in auditory sequences than non-musicians? If yes, is this advantage explained by an improved ability to learn sequence statistics or by an improved ability to process sensory information? Can we identify a neural correlate of the ability to learn sequence statistics? If yes, does this neural response possess an organized temporal structure? To answer these questions, we first evaluated whether musicians have better SL abilities than non-musicians by measuring their ability to predict the forthcoming items of auditory sequences that embed either simple or complex statistics. We then tested whether the advantage of musicians over non-musicians was explained by the ability to learn sequence statistics or by the ability to process sensory information by fitting a Bayesian model and contrasting individual parameters. We identified a neural correlate of the musician’s advantage by correlating the Bayesian model surprise with the EEG response amplitude at each time point, and compared the strength of this correlation in musicians and non-musicians. Finally, we explored the temporal structure of the EEG response by correlating three Bayesian models computing statistics at three levels of complexity with the EEG response amplitude at each time point.

## Results

### Musicians perform better than non-musicians in an auditory task implicating SL

We presented sequences of 300 items, drawn from a vocabulary of size 3 (impact sounds of glass, wood and metal ^19^). Two types of sequences that embed either simple or complex statistics were designed. Simple statistics corresponded to 1^st^ order Markov chains while complex statistics corresponded to 2^nd^ order Markov chains (see Figure 1A, see Methods). For each sequence, 20 probes were presented, asking participants to explicitly predict the most likely future item (glass, wood or metal). Note that the number of transition probabilities to be tracked grows exponentially with the order of the Markov chain, making complex sequences (P(⚪|⚪⚪), P(⚪|⚫⚪), …, P(⚫|⚫⚫) → 27 probabilities in total) harder to predict than simple ones (P(⚪|⚪), P(⚪|⚫), …, P(⚫|⚫) → 9 probabilities in total).

**Figure 1.**
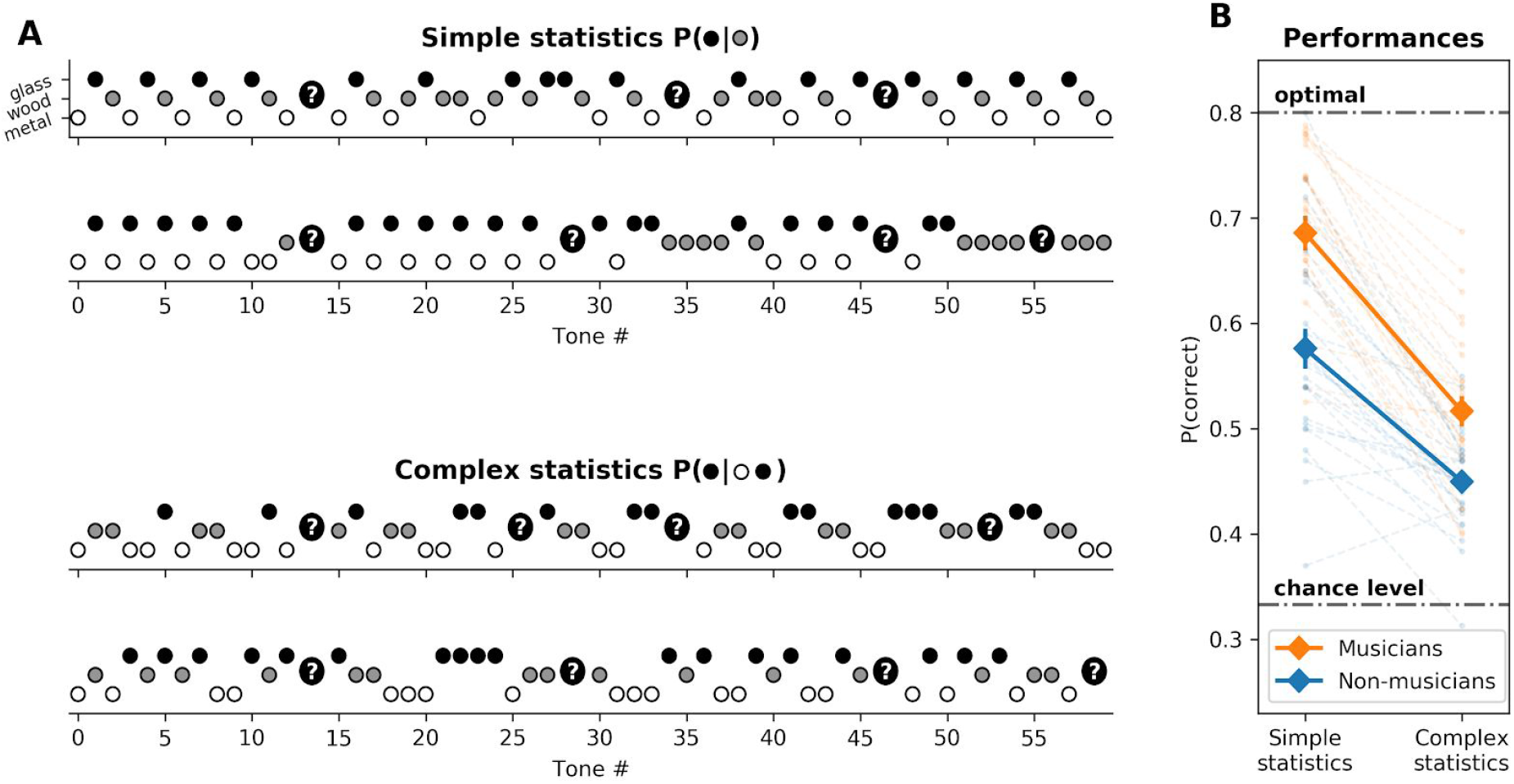
Musicians are better than non-musicians at predicting the forthcoming items of auditory sequences that embed either simple or complex statistics. **A.** Paradigm. Each sequence was composed of 300 tones, chosen among 3 (impact sounds of glass, wood and metal; here respectively black, grey and white). Two types of sequences were generated, two examples of each are shown: (top sequences) Simple statistics: 1^st^ order Markov chains, defined by P(s_t_|s_t−1_). Each item is chosen based on the previous one. (bottom sequences) Complex statistics: 2^nd^ order Markov chains, defined by P(s_t_|s_t−2_s_t−1_). Each item is chosen based on the previous pair of items. Participants were randomly probed for an explicit prediction about the forthcoming tone 20 times per sequence, here symbolized by question marks. **B.** Behavioral results. Overall, the performances are higher than chance (33%) but lower than the theoretical optimum (80% — sequences are probabilistic therefore 100% is not achievable). Participant’s predictions are closer to the generative statistics for simple sequences, compared to complex sequences (p < 10^−16^). On both types of sequences musicians are better than non-musicians (p < 10^−7^). Error bars represent standard error of the mean (s.e.m.).

Results were analyzed using mixed-effect logistic regression (see Figure 1B, see Methods). First of all, participants are better at predicting simple statistics (63.2 ± 1.5 %) than complex ones (48.4 ± 1.0 %, *β* = −0.51 ± 0.06, p < 10^−16^). In other words, while all participants perform well above chance level (33%), their predictions are closer to the generative statistics for simple sequences compared to complex sequences. It should be noted that performances are intermediate between a random strategy (33 %) and the theoretical optimum (80 % - sequences are probabilistic therefore 100% is not achievable). On average, musicians (60.1 ± 2.2 %) are better than non-musicians (51.3 ± 1.9 %), and this effect is significant (between-group comparison *β* = 0.48 ± 0.09, p < 10^−7^). This difference holds for both types of sequences. Musical expertise is associated with higher performances in a statistical learning task, for both simple (*β* = 0.48 ± 0.11, p < 10^−4^) and complex (*β* = 0.27 ± 0.07, p < 10^−3^) statistical regularities. Control analysis revealed that musicians and non-musicians do not benefit from an overall increase in performances during the course of the experiment (effect of block rank *β* = −0.001 ± 0.01, p = 0.84, interaction with musicianship *β* = 0.02 ± 0.01, p = 0.10), ruling out the possibility of a difference between groups due to task learning. Given the probabilistic nature of the task and the multiplicity of involved cognitive processes, the fact that musicians perform better than non-musicians can receive different explanations. We isolated four alternative hypotheses: (H1: memory span) musicians use a longer history of stimuli to make their predictions, (H2: sensory input quality) musicians are better at discriminating between sounds, (H3: SL) musicians estimate more complex statistics and (H4: selection noise) musicians have less noise in the selection process, *i.e.* they lose less information in late stages of the statistical learning process. The major problem is that these hypotheses make very similar predictions in terms of expected results: lower memory, increased sensory noise, inappropriate statistics and increased selection noise all provoke a decrease in average performances. Sticking to performance differences is therefore not the right approach to address this question. Instead, we relied on modelling tools: we designed a Bayesian model that makes explicit predictions about trial-by-trial responses and fitted it to the participants’ responses.

### Musicians estimate higher order transition probabilities (K) with a lower selection noise (*σ*) compared to non-musicians

The Bayesian model we designed (see Figure 2A, see Methods and Supp. Methods for a formal derivation of the model) is an “ideal observer” in the sense that it exploits all available information to compute predictions: it inverts the correct generative model, without noise and with perfect memory. Its predictions are encoded in a “transition probability matrix”, that links immediate contexts and items. This matrix is continuously updated using Bayes rule, given the observed sequence. The Bayesian model was fitted to each participant trial-by-trial responses (see Methods, see Figure 2B, Fig. Supp. 1). This procedure leads to a set of four parameters per participant: *λ*, *σ*, K, β. On average, the model explains 61.9 % (± 1.3) of the responses (musicians: 66.8 % ± 1.7; non-musicians: 56.8 % ± 1.5). It should be noted that this is higher than the performances of the participants (60.1 ± 2.2 %, p < 0.05), indicating that the model predicts the participants’ correct responses as well as their errors.

**Figure 2.**
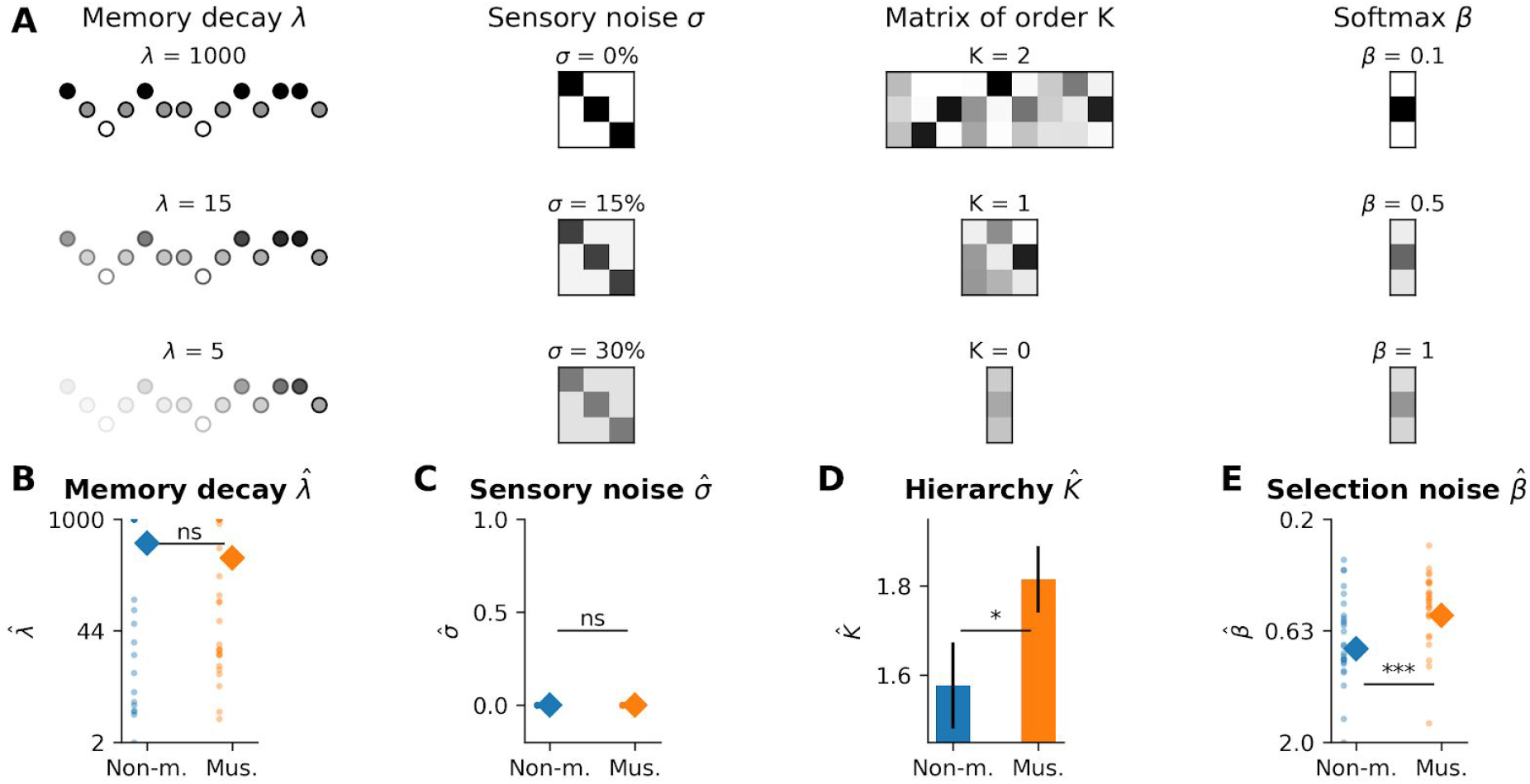
Bayesian modelling reveals key differences between musicians and non-musicians learning strategies. **A.** Bayesian model. The model comprises four components. (left) The sequence is weighted over time by an exponential decay function of parameter *λ*, mimicking memory forgetting (low values deteriorate performances). (middle left) The model receives inputs corrupted through a confusion matrix (deviation from the identity matrix deteriorates performances). (middle right) The model learns the transition probability matrix between contexts and items. The context can be composed of K = 1 or K = 2 items. The model can also be sensitive to the probability of an item without any reference context (K = 0). (K < 2 deteriorates performances for complex sequences). (right) The values of the transition probability matrix are converted into choice probability via a softmax rule, controlled by a selection noise parameter β (high values deteriorate performances). Musicians have **(B)** a non-significantly different memory decay parameter *λ* (p > 0.05), **(C)**, a non-significantly different sensory noise confusion matrix *σ* (p > 0.05) **(D)** a higher hierarchy parameter K, indicating a use of more complex statistics (p < 0.05) **(E)**, and a lower selection noise β (p < 10^−2^) compared to non-musicians.

The comparison of the fitted parameters revealed key differences between the two groups. (H1) Memory decay *λ* fitted values are similar between musicians and non-musicians (*β* = −199 ± 128, p = 0.13, ΔBIC = −1.5). (H2) The sensory noise is modelled as a “confusion matrix” and is also similar between musicians and non-musicians. For all participants, this best fitting matrix is the identity matrix, *i.e.* items are not confounded. This is coherent with the fact that the items were easily discriminable. Indeed, participants score 97.8 % (± 0.5) of correct responses in an item identification task during the familiarization block. (H3) The order K of the estimated transition probability matrix is higher in musicians compared to non-musicians (*β* = 1.39 ± 0.67, p < 0.05, ΔBIC = 0.78). This indicates that musicians tend to estimate more complex statistics. More specifically, the statistics they estimate (81 % of participants at K = 2) is the same as the statistics that generates the most complex sequences, *i.e.* 2^nd^ order Markov chains. Non-musicians apply simpler statistics (58 % of participants at K = 2) leading to a loss of performance compared to musicians. (H4) Finally, the selection noise *β* is lower in musicians compared to non-musicians (*β* = −0.26 ± 0.08, p < 10^−2^, ΔBIC = 6.3). Critically, the individual values of selection noise do not correlate with performances (*β* = −1.38 ± 1.15, p = 0.23) nor with reaction times (*β* = 0.21 ± 0.22, p = 0.37) in the item identification task of the familiarization block. This indicates that *β* does probably not represent response mapping confusion but rather genuine noise in the late stages of the statistical learning. Without being exhaustive, we can hypothesize for example greater computational precision, less over/underestimation of small/large probabilities or application of accurate heuristics that marginally approximate Bayesian computations.

### The P300 amplitude is more strongly correlated to Bayesian model surprise in musicians relative to non-musicians

We then used the Bayesian model fitted on the behavioral data to shed light on the brain responses. Following previous work ^14,20,21^, we hypothesized that brain signals linearly scale with the level of theoretical surprise. We relied on information theory ^22^ to formally define theoretical surprise as the negative log probability under the model M that, given a context, the forthcoming item will be a given item. This quantity corresponds to the intuitive notion of surprise: it is low when the item is expected and high when unexpected. We defined M as our Bayesian model fitted on behavioral responses. Formally, it was defined as −log_2_(P) where P is the posterior predictive probability of the presented item under the model M. We then associated a level of theoretical surprise with each item of each sequence for each participant. As only ~7% of items were behavioral probes, ~93% could be used to study EEG responses.

We relied on spatial filtering to reduce the dimension of the EEG dataset. Using multiway canonical correlation analysis ^23^, we computed spatial filters that maximize the temporal correlation between participants without diminishing inter-individual amplitude differences (see Methods, Fig. Supp. 2). The filters are summary components (SC), ordered by explained variance. The first SC was a central (frontal positive, occipital negative) filter, with a standard auditory response topology. It explained on average 55% of the variance of the ERP (peak correlation between SC time course and ERP at Fz, 93% explained variance) and was therefore kept for the rest of the analyses. We fitted a linear regression across all items between the theoretical surprise level and the SC amplitude at each time point. The matrix of linear coefficients (number of participants x number of time points) was then submitted to a cluster permutation in time algorithm to assess its statistical significance (see Figure 3A-B). The linear regression is significant during a long-lasting late time window for musicians (210 - 320ms, p < 10^−3^) and non-musicians (220 - 430ms, p < 10^−3^). The associated function, topography and time window of this response coincides with a P300 response. During active processing of the sequence, the amplitude of the P300 is therefore linearly strongly correlated with the theoretical level of surprise.

**Figure 3.**
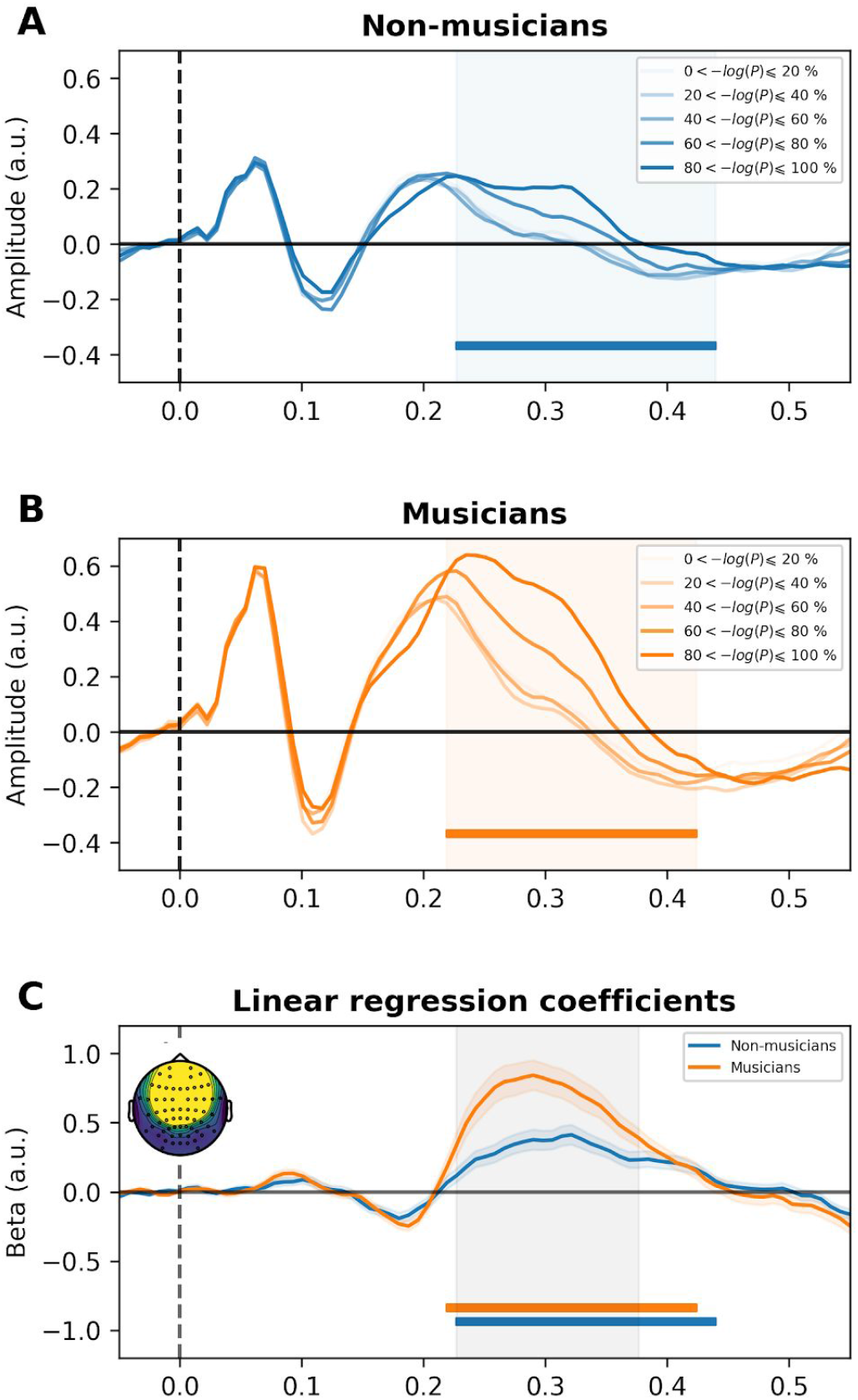
Bayesian agent surprise correlates with single trial late auditory event related potential (ERP) amplitude. **A.** Average ERP of the main summary component (SC, see Methods) of non-musicians as a function of their model surprise quintiles. Model surprise was defined as −log_2_(P) where P is the posterior predictive probability of the presented item under the Bayesian model fitted to the participants behavioral responses. The linear regression is significant in a late time window, between 220 and 430 ms. **B.** Average ERP of the main SC of musicians as a function of their model surprise quintiles. Significant time cluster is shown in orange. The linear regression is significant in a late time window, between 210 and 420 ms. **C.** Coefficients of the linear regression between model surprise and SC amplitude for musicians and non-musicians. The difference is significant in the same late time window, between 220 and 380 ms. Significance of the effect (all p < 10^−3^) was assessed using correction at the level of the cluster (blue and orange lines). Significance of the difference between groups (p < 0.05) was also cluster-corrected (grey area). Colored shaded areas represent standard error of the mean (s.e.m.). Inset plot shows the main SC topography.

**Figure 4.**
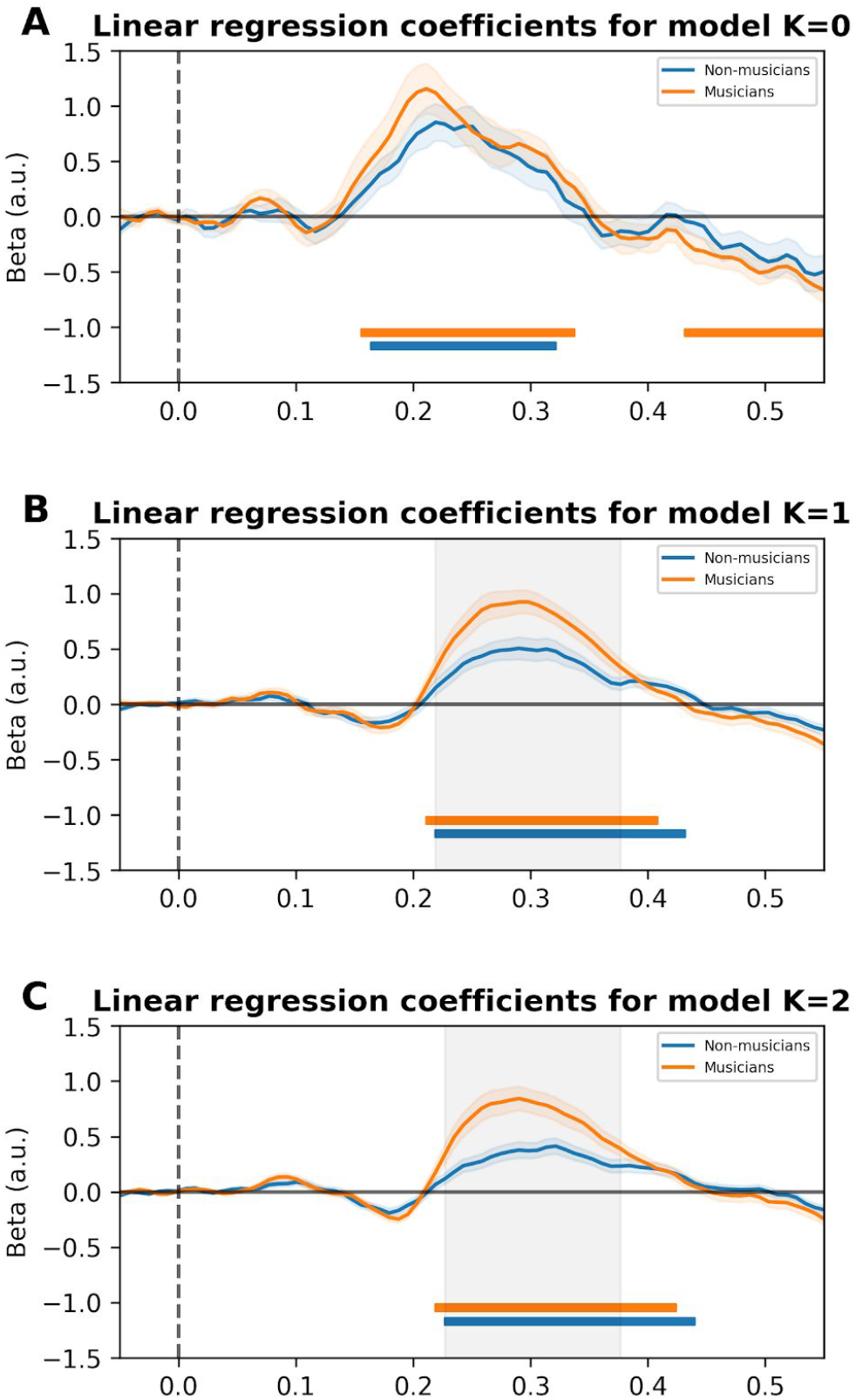
A cascade of low to high complexity surprise correlates with the amplitude of single trial auditory event related potentials (ERP). **A.** Coefficients of the linear regression between surprise from the simplest model (K=0) and SC amplitude for musicians and non-musicians. The linear regression is significant in a middle latency window, between 150 and 300 ms. There is no significant difference between groups (p > 0.05, cluster-corrected) **B.** Coefficients of the linear regression between the model (K=1) surprise and SC amplitude for musicians and non-musicians. The linear regression is significant in a late latency window, between 210 and 430 ms. The difference between the groups is significant (p < 0.05) in this late time window, between 220 and 380 ms (shown in grey). **C.** Coefficients of the linear regression between the most complex model (K = 2) surprise and SC amplitude for musicians and non-musicians. The linear regression is significant in a late latency window, between 210 and 430 ms. The difference between the groups is significant (p < 0.05) in this late time window, between 220 and 380 ms (shown in grey). Significance of the linear regression is shown in orange and blue (all p < 10^−3^). Colored shaded areas represent standard error of the mean (s.e.m.).

We finally submitted the difference between musicians and non-musicians coefficient matrices to a cluster permutation algorithm. This analysis revealed that the linear coefficients of the musicians are higher than the coefficients of the non-musicians in the same late time window (220 - 380ms, p < 0.05). Using an unbiased estimator such as the coefficient of determination R^2^ instead of the linear coefficients leads to similar results (see Fig. Supp. 3). This ensures that the difference is not explained by the difference of overall ERP amplitude between musicians and non-musicians. The P300 amplitude is therefore modulated by the theoretical surprise to a higher degree in musicians than in non-musicians (see Figure 3C).

### The ERP amplitude modulation enhancement in musicians is restricted to complex statistical learning

The key component of the model is the estimation of the transition probability matrix. Critically, this matrix can be of any order (K), *i.e.* reflects the probability of observing an element given the preceding item (K = 1), bigram (K = 2), trigram (K = 3), or even *a priori*, without context (K = 0). The order of the matrix allows to order SL complexity, from simple (K = 0) to complex predictions (K ≥ 1). The idea of ordering is necessary in the context of hierarchical theories of brain organisation ^24–26^, in particular the Bayesian brain hypothesis. The Bayesian brain hypothesis states that the brain is a hierarchical dynamical system composed of nodes, whose activity reflects the difference between predictions coming from top (associative) nodes and prediction errors coming from bottom (sensory) nodes. The overall computational goal of the system is to build an appropriate internal model of the world, in order to predict its future states. In this view, our task would first prompt simple statistical estimates (K = 0), whose activity will then be regulated by more complex predictions that take immediate context in account (K ≥ 1).

To test this hypothesis, we relied on the Bayesian model. We fixed the previously fitted parameters except one: the order parameter K. This allowed us to define a level of theoretical surprise to each item, each sequence, each participant, for simple (K = 0), medium (K = 1) and complex statistics (K = 2). We then performed the same linear regression followed by cluster permutation analysis. The regression with simple statistics (K = 0) showed a modulation peaking around 200 ms, with a non-significant tendency for an earlier latency and greater amplitude for musicians compared to nonmusicians. Contrast analysis revealed that the regression coefficients are higher for simple statistics (K = 0) than for more complex ones (K = 1 and K = 2) in the same time window (both p < 0.05). The topology and time window of the modulation is consistent with a MMN ^27^. This is coherent with the fact that a “K = 0” model is actually estimating the probability of an item irrespective of its immediately preceding context, *i.e.* the overall frequency of occurrence of this element. Indeed, MMN are typically elicited in an oddball paradigm, wherein the frequency of occurrence of items is manipulated (frequent *versus* rare). By contrast, the regression with complex statistics revealed a significant correlation with the ERP amplitude later in time, between ~200 and ~400 ms for both “K = 1” and “K = 2” models. Group contrast analysis revealed that this modulation is larger for musicians compared to non musicians for both higher level models (p < 0.05). Contrast analysis revealed that the regression coefficients are higher for complex statistics (K = 1 and K = 2) than for simple ones (K = 0) in the same time window (both p < 0.05). Using the coefficient of determination R^2^ instead of the linear coefficients leads to similar results (see Fig. Supp. 4). In a nutshell, the simple statistics model correlates with ERP amplitude around 200 ms, similar to an MMN, and this correlation is similar in musicians and non-musicians. By contrast, more complex models correlate around 300 ms, similar to a P300, and this correlation is higher in musicians compared to non-musicians.

## Discussion

Musicians make more accurate predictions than non-musicians in a complex auditory SL task. Computational modelling reveals that this advantage is best explained in terms of parameters governing the ability to process SL (order of the Markov chain model and selection noise), as opposed to parameters governing the ability to process sensory information (sensory input noise, memory decay). EEG recordings during behaviorally unprobed items allow bridging modeling and electrophysiological signatures of the behavioral task. First, the amplitude of a central frontal cluster at 300 ms is strongly correlated on a single trial basis to the computationally modeled theoretical surprise. Second, this P300 model-based modulation is stronger for musicians than non-musicians, suggesting a difference in sensitivity to the probabilistic structure of the sequence. Last, musicians and non-musicians have similar neural responses to surprise in a simple statistical structure (K = 0), and diverge only for surprise in more complex statistical structures (K = 1 and K = 2). We conclude from these results that musicians have improved neural SL compared to non-musicians.

Our results shed light on the musical training induced plasticity. It has been known for a long time that musical training is associated with low level sensory improvements, such as pitch or duration detection thresholds ^28–30^. However, music expertise does not only require fine perception of isolated pitches and durations but it also puts high demands in terms of sequence processing (melody and harmony) that require both accurate temporal and spectral prediction ^31^. In the current experimental design, we chose to control sensory-related gain biases by using large acoustic differences between stimuli. Thus, it is not surprising that the model does not capture low level sensory differences. By contrast and critically, the model captures differences related to downstream computations, directly involved in the inference process itself. It is important to note that the Bayesian model not only captures the surprise associated with violations of the internal expectations but it also describes the continuous learning by constantly updating predictions as a function of the context. Hence, the advantage of musicians over non-musicians does not solely rely on a greater sensitivity to predictions errors, as shown by the larger P300 modulation, it also reflects a more complex and more accurate continuous learning. This advantage is manifest in the hierarchy of ERP modulations, where modulations associated with simple statistics prediction (K = 0) do not differ between groups while more complex statistics prediction modulations do differ (K ≥ 1). Our results replicate and integrate in the Bayesian framework the notion of “abstract” MMN ^32^, *i.e.* the sensitivity to regularities beyond the mere frequency of appearance. Studies have demonstrated that musicians have larger MMN in response to changes in melodic contours ^33–36^, while having similar MMN in response to simple change of pitch ^36^. Similar findings have also been described for chord sequences ^37^. Interestingly, changes of pitch concern the frequency of occurrence of items, *i.e.* K = 0 Markov chains, whereas changes of melody concerns the ordered structure, which is typically captured by a K = 1 or K = 2 Markov chain. We extend these results by using a task that does not rely on the pitch dimension, thus ensuring that sensory processing is not the origin of the musical expertise advantage. Furthermore, the Bayesian model we developed sheds light on these results by drawing an explicit line between “simple” statistics (frequency of occurrence, K = 0) and “complex” ones (K ≥ 1). Anecdotally, the musical advantage of complex statistics over simple ones is also present in music automatic generation, where high order Markov chains “generate results with a sense of phrasal structure, rather than the ‘aimless wandering’ produced by a first-order system” ^38^ (see Iannis Xenakis’ Analogique A and B for an example music generated by 1st order Markov chains).

Our Bayesian model combines two approaches: an “ideal observer” and an “individualized modelling” strategy. This ideal observer allows defining a theoretical maximum on the probabilistic task, here defining performance bounds between a random (33%) and an optimal strategy (80%). However, ideal observer models are usually universal, and as such are unable to characterize inter-individual variability. By contrast, the added parameters included in our model precisely specify the multiple ways of being suboptimal and allow testing clear and separable hypotheses about different suboptimality sources. Using this combined approach, we reveal that humans do not perform optimal predictions. If (Bayesian) optimality has been standard in psychophysics, *e.g.* in visual orientation discrimination ^39^ or multisensory integration ^40^ tasks, suboptimality is almost systematically observed in higher level cognitive task, such as discrete evidence accumulation ^41^ or explicit manipulation of probabilities ^42^. The errors are usually explained in terms of task-independent noise in sensory processing ^43–45^ or noise in the response selection, following the decision ^46^. However, recent proposals have put emphasis on limitations in the inference process itself ^41,47,48^, arising from systematic biases ^49^ or from variability in the computational precision of variables represented in populations of neurons ^50^. Our results confirm these recent proposals. First, the key parameters to separate musicians and non-musicians are not the sensory sources of noise but the complexity of the estimated statistics (K) and the selection noise (*σ*) after this computation. These results can directly be interpreted in terms of simpler and less accurately estimated statistics by non-musicians compared to musicians. Second, the ERP component sensitive to model surprise peaks rather late, around 300 ms post-stimulus onset, which is also indicative of a downstream process, as opposed to an early one, *e.g.* P1 or N1. Last, we also demonstrate that the complexity of the probabilistic structure affects the timing of neural computations in a hierarchical manner: simple statistics estimation rests on early neural modulations, around 200 ms, while more complex statistics estimation rests on late neural modulations, around 300 ms. Further work is necessary to precise the errors in the inference process. Indeed, although the complexity of the estimated transition probability matrix seems to be a good proxy of the difference between musicians and non-musicians, the neural mechanisms underlying the observed suboptimality are open to debate ^49,50^.

The hierarchy of EEG modulations that we observe is to be interpreted in the context of hierarchical theories of brain organisation, in particular the Bayesian brain hypothesis ^24,25^. In this view, sensory information is evidenced to be confronted with an internal model and its predictions. In EEG, the prediction errors typically induce an increase of P300 amplitude ^13,14,20,21^. In this experiment, we have operationalized the notion of hierarchy through the notion of order of a Markov chain, in the sense that higher in the hierarchy corresponds to higher in the order of the Markov chain. It should be noted that under the law of total probabilities, high order probability embed lower order ones, *e.g.* P(⚫) = P(⚫|⚪)P(⚪) + P(⚫|⚫)P(⚫) - in a universe where only ⚫ or ⚪ can happen. This essentially means that a node that is computing complex statistics needs to receive input from a node that is computing simple ones. In particular, it must receive its prediction errors in order to update its values. This simple law imposes a temporal ordering on the predictions errors: we should observe ERP modulation in an organized manner, from simple (K = 0) to complex (K = 2) prediction errors. This is precisely what we show. Furthermore, the Bayesian framework also offers a mechanistic explanation of the impact of musical training on the brain. By providing the right amount of predictability and prediction errors at different levels of complexity ^51^, music training would shape brain networks in order to facilitate hierarchical predictions.

Finally, as a fundamental computation for any temporal structure representation, SL is a major target for multiple cognitive functions rehabilitation: for language by promoting word segmentation ^5^, gesture learning by enhancing motor sequence learning ^52^ or working memory by allowing chunking strategies ^53^. Theories of learning, like the reverse hierarchy theory ^26,54^, state that learning is more effective in high order processing than in low order ones. We demonstrate here that inter-individual differences linked to musical training do not concern low level abilities, thus providing encouraging results for rehabilitation. As an extensive complex SL training, musical training could be a very efficient way to remediate to a vast repertoire of cognitive functions. Caution should nonetheless be taken, as our results are of correlational nature ^55^. Although unlikely, it could be the case good SL abilities is actually a prerequisite for musical training, leading to a greater proportion of good SL performers in musicians compared to non-musicians. However, causal studies have already demonstrated that musical training is effective for speech segmentation based on SL ^56^, for rehabilitation of reading and phonological skills ^57^ in children with dyslexia, turn taking in children with cochlear implant ^58,59^ as well as in several neurological disorders (for a review, see ^60^). Since temporal structuring is most often impaired, we formulate the mechanistic hypothesis that SL rehabilitation may be at the core of the music remediation power. By providing a probabilistic temporal structure and focusing attention on this structure, music could lead to beneficial effects on multiple cognitive abilities *via* SL training.

## Methods

### STIMULI AND PARADIGM

#### Participants

We collected data from 27 musician participants (17 females, mean age 33.3 y, standard deviation ± 12.2, range [18, 62]) and 26 non-musician participants (15 females, 31.1 y ± 11.3 [20, 55]) following informed consent. All had normal hearing, reported no neurological deficits and received 20 euros for their time. The experiment was approved by the National Ethics Committee on research on human subjects.

#### Stimuli

Across the experiment, 10 sequences of 300 items (sounds) were presented to the participants. The vocabulary consisted of three items A, B and C. Each sequence contained the same amount of item types (100 As, Bs and Cs). The order of the items of the 10 sequences was designed to probe two levels of statistical learning complexity. Five sequences were 1^st^ order Markov chains: each item was chosen only based on the previous item given the corresponding column in a transition probability matrix of size 3×3. This matrix described the probability of choosing a particular item given the preceding one, *e.g.* P(A|B). The matrix was biased so that each item was followed primarily by one item (p=0.8) compared to the other two (p=0.1). The other five sequences were 2^nd^ order Markov chains: each item was chosen only based on the previous pair of items given the corresponding column in a transition probability matrix of size 9×3. This matrix described the probability of a particular item given the preceding pair, *e.g.* P(A|CB). The matrix was biased so that each pair was followed primarily by one item (p=0.8) compared to the other two (p=0.1).

The three items A, B, and C were randomly assigned to three sounds for each participant. The sounds were artificially generated impact sounds : wood, metal and glass ^19^. Importantly, all sounds had the same fundamental frequency, loudness and duration, and differed only in timbre (examples of “tuned” sounds available at http://www.lma.cnrs-mrs.fr/~kronland/Categorization/sounds.html). Each sound was 150 ms long, with cosine ramp on and off of 10 ms, presented at a fixed rate of ~1.6 Hz (onset asynchrony of 600 ms). Each sequence lasted ~4.5 min.

Explicit prediction judgments were probed 20 times per sequence. Probes were randomly spaced by 9, 12, 15, 18 or 21 items. During a probe, participants had 4s of silent intervals to indicate the most likely forthcoming item, using key press on a keyboard.

#### Procedure

Classical SL paradigms are usually composed of two phases ^5,6,52^: a habituation block, consisting in the presentation of a sequence of items embedding the statistical regularity to be learned, and an evaluation block, consisting in the presentation of sequences that respect the regularity (“standard”) or not (“deviant”). Learning is indirectly measured as the impact of the violation of the rule, typically differences in reaction time or accuracy between “standard” and “deviant” items. Our paradigm differed in two aspects. (1) We evaluated SL during the learning block and suppressed the evaluation block. Beyond considerably reducing the duration of the experiment, this solves critical problems related to forgetting and learning occurring in the evaluation block ^9^. As the stimuli were probabilistic sequences and as learning is a continuous process, there was no *a priori* way to classify items into two binary classes, like “standard” or “deviant”. Instead, we relied on modelling tools and information theory to define the degree of expectation violation/fulfillment as the “theoretical surprise” elicited by each item. (2) The task of the participant was to make explicit predictions, therefore SL was not indirectly measured *via* infrequent violations of the rule, but directly *via* the accuracy of the predictions and the adequation between participants’ responses and model predictions.

Participants were seated in a soundproof room in front of a computer screen, a loudspeaker and a keyboard. Prior to the experiment, correct audition was assessed using a rapid 5dB-step audiogram. Participants were then familiarized with the stimuli and the mapping between the items and response keys in a block of 50 items. During this block, items were presented in a random order and participants had to press the key corresponding to the heard sound at every item (three keys “left”, “up” and “down”). The mapping between keys and sounds were fixed during the whole experiment and randomized across participants. On average, participants scored 97.8 % (± 0.5) of correct responses during this familiarization block.

Participants were then instructed to listen to the sequences, and to predict the forthcoming item using the keyboard whenever they saw the probe screen. Emphasis was put on accuracy and not speed. Each participant did 10 sequences, 5 of each type (1^st^ order and 2^nd^ order Markov chains), in a random order. Participants were aware of the probabilistic nature of the sequences and of the difference between the two types of sequences. They could take small breaks between each sequence. The whole experiment lasted ~50 min.

The sequences were presented binaurally to participants at an adjusted comfortable level (~70 dB) using loudspeakers. Stimuli presentation and data collection was controlled with Python custom scripts. Submillisecond synchrony between stimulus presentation and EEG acquisition was ensured using triggers embedded in the audio files and delivered to the acquisition computer via a dedicated channel.

### BEHAVIORAL ANALYSES

#### Outliers

One participant was removed from the analyses because of poor performances in the familiarization block (80%, < −3 on the z-scored performances scale of the group).

#### Statistical analyses

Statistical analyses were done using R and the package lme4 ^61^. The effects of sequence type, musicianship and their interaction on performances were estimated using logistic mixed-effect models. The probability of a correct response (0: incorrect, 1: correct) was modeled as a logistic regression with sequence type (0: 1^st^ order Markov chain, 1: 2^nd^ order Markov chain), musicianship (0: non-musicians, 1: musicians) and their interaction as predictors. Model complexity was monitored using the Akaike Information Criterion, a standard measure to arbitrate between complexity and accuracy. The most parsimonious model was a model including sequence type, musicianship and their interaction as fixed effects and the intercept of each participant as random effect. Reported p-values are Satterthwaite approximations.

### MODELING ANALYSES

#### Bayesian optimal model

The optimal model was based on a previously published model IDyOM ^62^, an n-gram model that we reframed and extended in a Bayesian framework. The goal of the model is to infer the probability of each item given the preceding context. Formally, the model is exposed to a sequence of t items s_0:t−1_ of vocabulary V of size 3. The context is given by the last K items of the sequence. As a Bayesian ideal observer, she uses Bayes rule to update her belief:

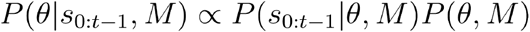

The transition probability matrix *θ* describes how likely each element is, given its preceding context. This learning process consists in estimating *θ* from the sequence s_0:t−1_. The full derivation of the likelihood term is given in Supplementary Methods. In the end, the predictive posterior probability of the model for each item is, rather naturally:

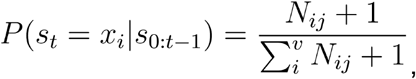

where P(s_t_ = x_i_|s_0:t−1_) is the probability of the element x_i_ ∈ V. It corresponds to the ratio of two scalars:

- N_ij_+1: the number of times that the context j and the element i have been observed together plus one,
- ∑_i_ (N_ij_+1): the number of times that the context j has been observed plus v.

#### Bayesian model

Suboptimality was introduced by four means to tear apart four hypotheses. First, the size of the context K varied between 0 and 2. As the generated sequences are Markov chains of order 1 and 2, only a model with K = 2 can properly do the task. Second, a leak parameter was introduced to account for memory decay. Previous observations were weighted by a weight e^−t/*λ*^ for *t*-th past stimulus. A small *λ* indicates quicker memory decay and worsen the performance. Finally, predictions were transformed into choice probability via a softmax rule with inverse temperature β. A high β indicates high noise in the decision process and worsens the performance.

#### Model parameter recovery

Before performing the experiment, we selected sequences that allow proper model estimation. We assessed our model selection procedure with a parameter recovery analysis. We generated synthetic data for 10^3^ models with random parameters (*λ* uniform between 2 and 1000, *σ*, exponential between 0 and 1, K, uniform choice between 0, 1 and 2, β exponential between 0 and 1). We then ensured that the recovered parameters from these synthetic data using our procedure were close to the original parameters. This was the case using our sequence and the same number of trials as our participants (spearman *ϱ*_*λ*_ = 0.78, *ϱ*_*σ*_ = 0.62, *ϱ*_K_= 0.90, *ϱ*_β_ = 0.92, see Fig. Supp. 5).

#### Model fitting

The model fitting values reported in the paper are maximum likelihood estimates (see Fig. Supp. 1). Formally, for a set of responses r_0:N_, assuming that each response is independent, the likelihood was defined for each participant as:

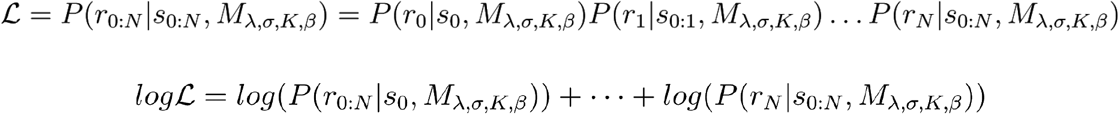

As the logarithm is a monotonically increasing function, finding the maximum of the likelihood is the same as finding the maximum of the log-likelihood. As the number of parameters is low (*λ*, *σ*, K, β), we relied on grid search to find estimates of this maximum. We computed the log-likelihood for each participant on trial-by-trial responses, for 200 logarithmically spaced values of *λ* (range 1, 1000), 200 logarithmically spaced values of *σ* (range 0, 2), 3 values of K (range 0, 2), and 200 logarithmically spaced values of β (range 0, 2). For each participant, the argmax of the log-likelihood on this grid was defined as its parameters estimates.

### ELECTROENCEPHALOGRAPHY (EEG)

#### Apparatus

EEG signal was recorded at 1000 Hz sampling rate using a BrainAmp amplifier and 64 preamplified Ag–AgCl electrodes mounted following the 10–10 international system (actiCap) in a soundproof and Faraday cage. The ground electrode was placed at AFz and the reference electrode at FCz.

#### Preprocessing

Signal processing was done using MNE-python ^63^ and custom scripts written in Python. Continuous data were bandpass filtered (1-40 Hz, zero-phase Hamming window FIR filter) and major artifacts rejected by visual inspection. Independent component analysis (fastICA) was used to remove physiological artifacts such as eye blinks and muscular activity. Data were then segmented into epochs of 600 ms starting at 50 ms prior to item onset and stopping 550 ms after. Epochs were zero-mean normalized to baseline ([−50, 0] ms) and re-referenced to the algebraic average of all electrodes.

#### Multiway canonical correlation analysis (MCCA)

Brain signals recorded with EEG have poor signal-to-noise ratio due to the presence of multiple competing sources and artifacts. A common remedy is to average recorded signals over multiple repetitions of the same stimulus and over multiple participants. However, averaging across participants is problematic, because differences in brain sources and geometry considerably increase variance. To deal with this problem, we relied on a powerful yet simple method recently developed ^23^. Multiway canonical correlation analysis (MCCA) consists of summarizing the data into individual spatial filters, named “summary components” (SC). These individual filters are built to maximize the temporal correlation between participants. MCCA was run on pooled data of musicians and non-musicians, in order to avoid spurious group differences. The first SC explained on average 55% of the variance of the ERP (peak correlation between SC time course and ERP at Fz, 93% explained variance) and was therefore selected for the rest of the study. Electrodes best explained by the SC were FC1, FC2, F1, Fz, F2, FC4, FC3, C2, C1, F4, F3, Cz, C3, which is consistent with an auditory response topography.

#### Model surprise regression

As the stimuli are probabilistic sequences and as learning is a continuous process, there is no *a priori* way to classify items into two binary classes, like “expected” or “unexpected”. Instead, we relied on information theory ^22^ to define the degree of expectation as the “surprise” elicited by each item. Formally, the surprise is defined as −log_2_P(x_i_) where P(x_i_) is the posterior probability P(s_t_ = x_i_|s_0:t−1_, M_*λ*,*σ*,K,β_) of the model M_*λ*,*σ*,K,β_ on the presented item s_t_. Critically, the surprise depends on a particular model of the world M_*λ*,*σ*,K,β_, that ascribes a probability P(s_t_ = x_i_) to each possible item at each time step t. We defined M_*λ*,*σ*,K,β_ as our Bayesian model fitted on behavioral responses. We then associated a level of theoretical surprise to each item of each sequence for each participant.

We therefore used the parameters extracted by model fitting from the behavioral responses (that concern only a small fraction of all items) to study the EEG responses to each item (280 items x 10 sequences per participant). A value of surprise was defined for each participant, each sequence, and each item, based on the behaviourally fitted parameters. We regressed the surprise against the SC amplitude in epochs that did not correspond to behavioral probes (280 items x 10 sequences). For that, we computed the linear regression of the surprise at each time point, ending up with an array of linear coefficients of size 52 × 600.

Statistical significance of the coefficients was assessed using cluster permutations in time (n = 2048). A t-test against 0 was performed and cluster-corrected for each group to assess the significance of the linear regression. An independent t-test was performed and cluster-corrected between groups to assess the significance of the difference of the coefficients between musicians and non-musicians.

## Acknowledgments

We thank Céline Hidalgo and Patrick Marquis for helping with the data acquisition, Clement François, Benjamin Morillon, Maxime Maheu and Christopher Summerfield for very helpful comments on previous versions of the manuscript.

## Supplementary Methods

### DERIVATION OF THE BAYESIAN OPTIMAL MODEL

The goal of the model is to infer the probability of each item given the preceding context. Formally, the model is exposed to a sequence of t items s_0:t−1_ of vocabulary V of size v. The context is given by the last K items of the sequence. As a Bayesian ideal observer, she uses Bayes rule to update her belief:

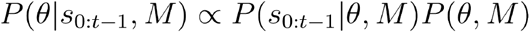

We can decompose the likelihood term using the chain rule :

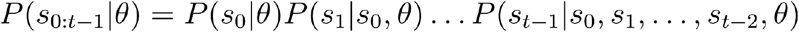

#### Restricted case K = 1, v = 2

For simplicity, let’s first suppose a restricted case in which K = 1, *i.e.* the sequence is a Markov chain of order 1, and v = 2, *i.e.* there are only two items: A and B. The derivation is inspired from ^15,64^. The likelihood of a given observation depends only on the estimated transition probabilities and the previous item:

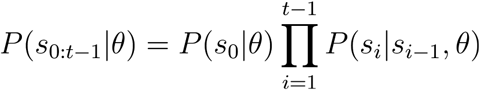

Assuming that for the first observation P(s_0_|*θ*) = 0.5, we have:

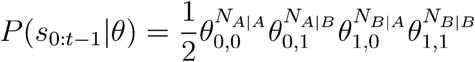

Where *θ* denotes the “transition probability matrix” of order 1. It is a matrix of size 2 × 2 where each row designates a particular item (A or B) and each column a particular context item (A or B). N_A|B_ designates the number of occurence of the bigram “BA” in the sequence, *i.e.* the number of times that the context “B” was followed by the item “A”. As each column sum to one, we can rewrite the equation as:

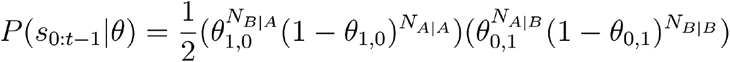

A nice conjugate prior for this likelihood equation is the Beta distribution as the product of two Beta distributions is also a Beta distribution. Combined with a uniform prior distribution Beta(1, 1), the posterior distribution on *θ* is therefore the product of two Beta distributions with parameters corresponding to the transition counts plus one.

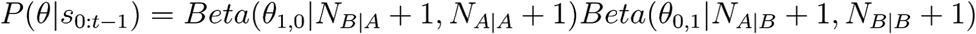

#### General case

The general case is similar to the restricted case. The main extension relies on the fact that the Dirichlet distribution generalizes the Beta distribution, with more than two parameters. The only difference will therefore be that the posterior is not any more a product of two Beta distributions but a product of v^k^ Dirichlet distributions with v parameters. As previously, the likelihood of a given observation depends only on the estimated transition probabilities and the previous K items:

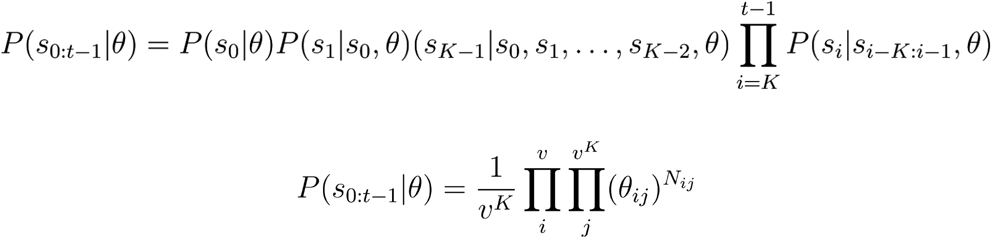

Where *θ* denotes the “transition probability matrix” of order K. It is a matrix of size v × v^K^ where each row designates a particular item and each column a particular K-uplets of items. N_ij_ designates the number of occurrences of the (K+1)-uplet corresponding to the cell (i, j) of the matrix (the j-th K-uplet followed by the i-th item). For simplicity, the first K observations are considered arbitrary such that P(s_0_) = P(s_1_) = … = P(s_K–1_) = 1/v. As each column sums to one, the derived likelihood corresponds to the product of v^k^ Dirichlet distributions. Combined with a uniform joint prior distribution Dir(1, 1, …, 1), the posterior distribution therefore results in a Dirichlet distribution with parameters corresponding to the transition counts N_ij_ plus one:

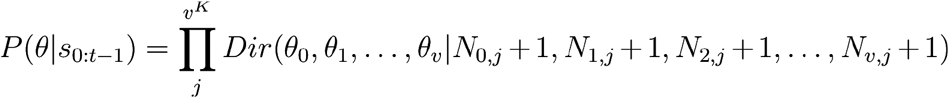

The posterior distribution can then be turned into the likelihood of the next stimulus using Bayes’ rule:

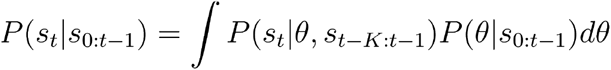

Which can be analytically solved and and ends up being simply:

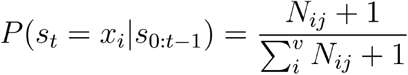

Where P(s_t_ = x_i_|s_0:t−1_) is the probability of the element x_i_ ∈ V. It corresponds to the ratio of two scalars:

- N_ij_+1: the number of times the (K+1)-uplet s_t−K_s_t−K+1_…s_t−1_x_i_ has been observed plus one,
- ∑_i_ (N_ij_+1): the number of times that the K-uplet context s_t−K_s_t−K+1_…s_t−1_ has been observed plus v.

## Supplementary Figures

**Figure Supp. 1.**
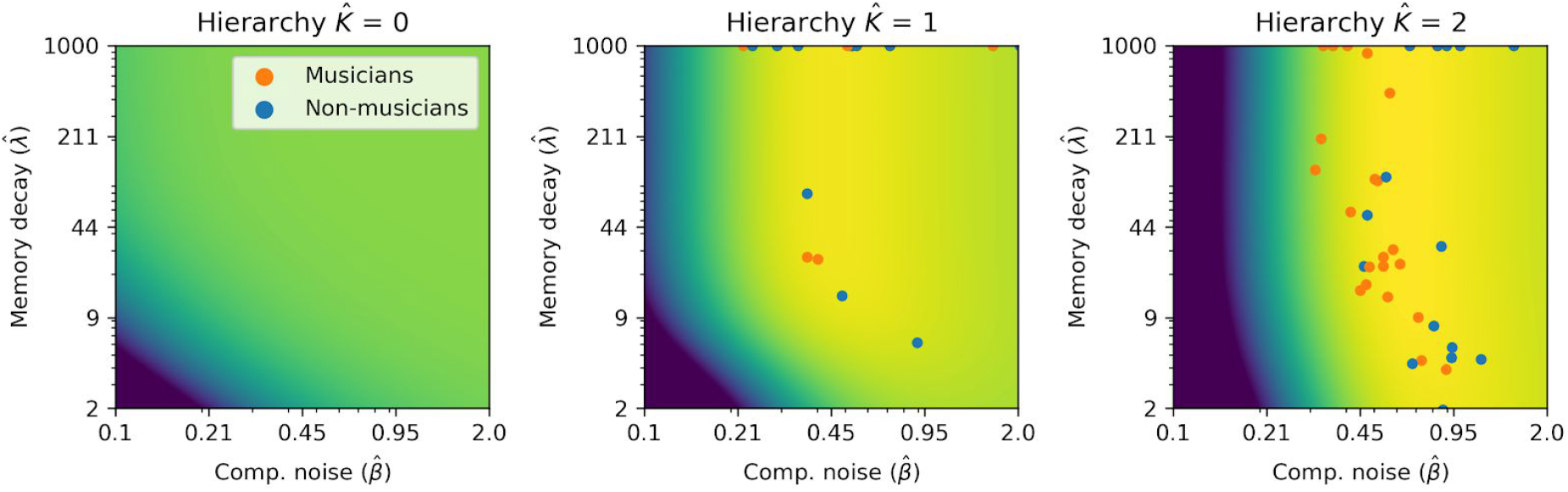
Maximum-likelihood fitting. Average log-likelihood of the data as a function of model parameters. Yellow indicates higher log-likelihood. Colored dots represent individual participants. The fitting of *σ* have been omitted for visualization purpose ― its value was 0 for every participant.

**Figure Supp. 2.**
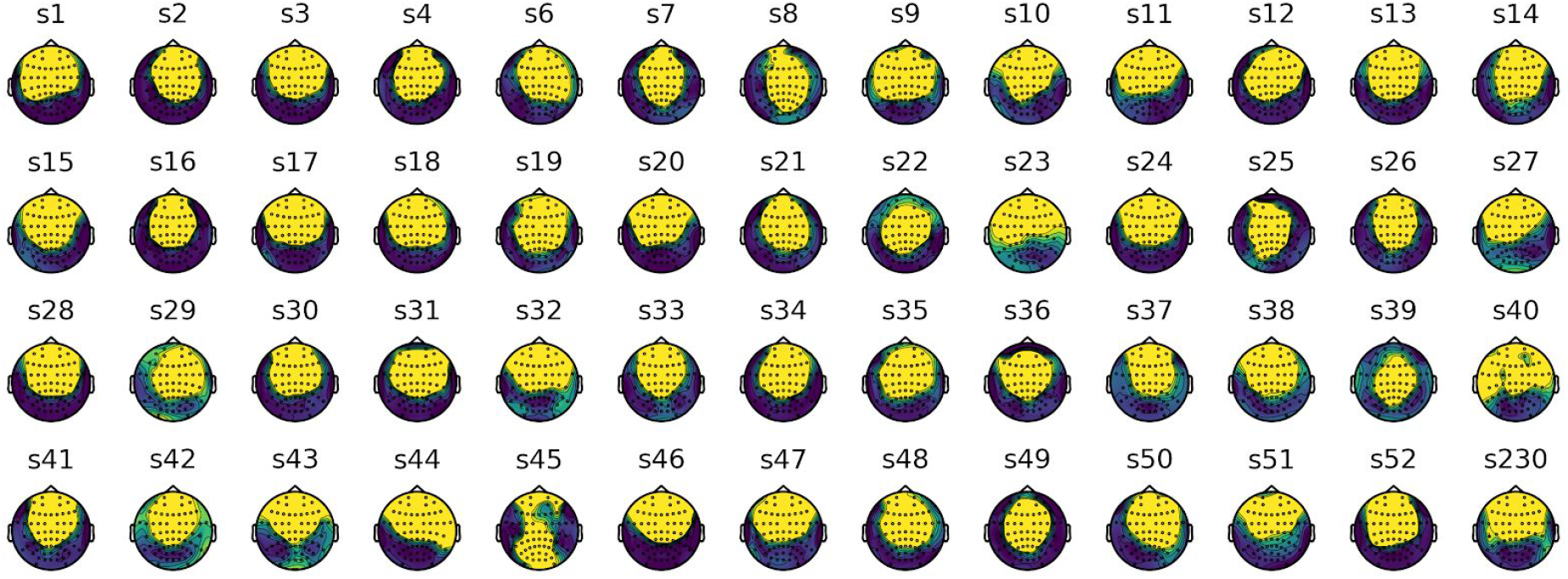
Topographies of the Summary Component (SC) #1 computed by the Multiway Canonical Correlation Analysis. Each topography represents the correlation between one individual SC #1 time course and each electrode time course. High correlation (yellow) indicates that the electrodes contributes strongly to the SC #1 time course.

**Figure Supp. 3.**
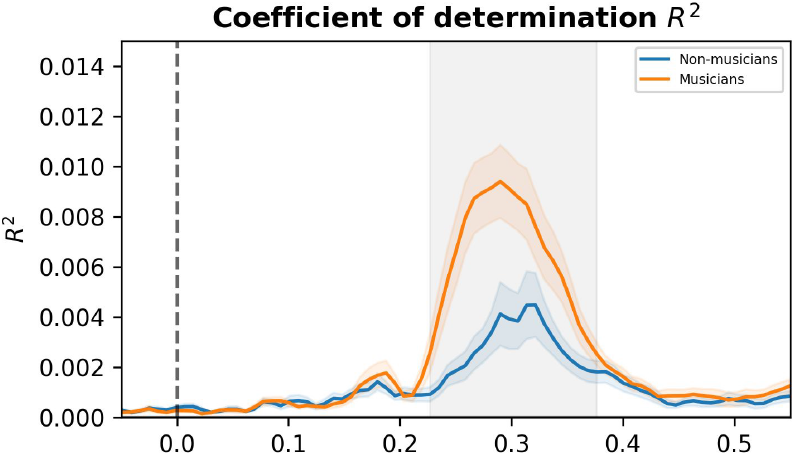
Coefficient of determination R^2^ of the linear regression between model surprise and SC amplitude for musicians and non-musicians. The difference is significant in a late time window, between 220 and 380 ms. Significance of the difference between groups (p < 0.05) was cluster-corrected (grey area). Colored shaded areas represent standard error of the mean (s.e.m.).

**Figure Supp. 4.**
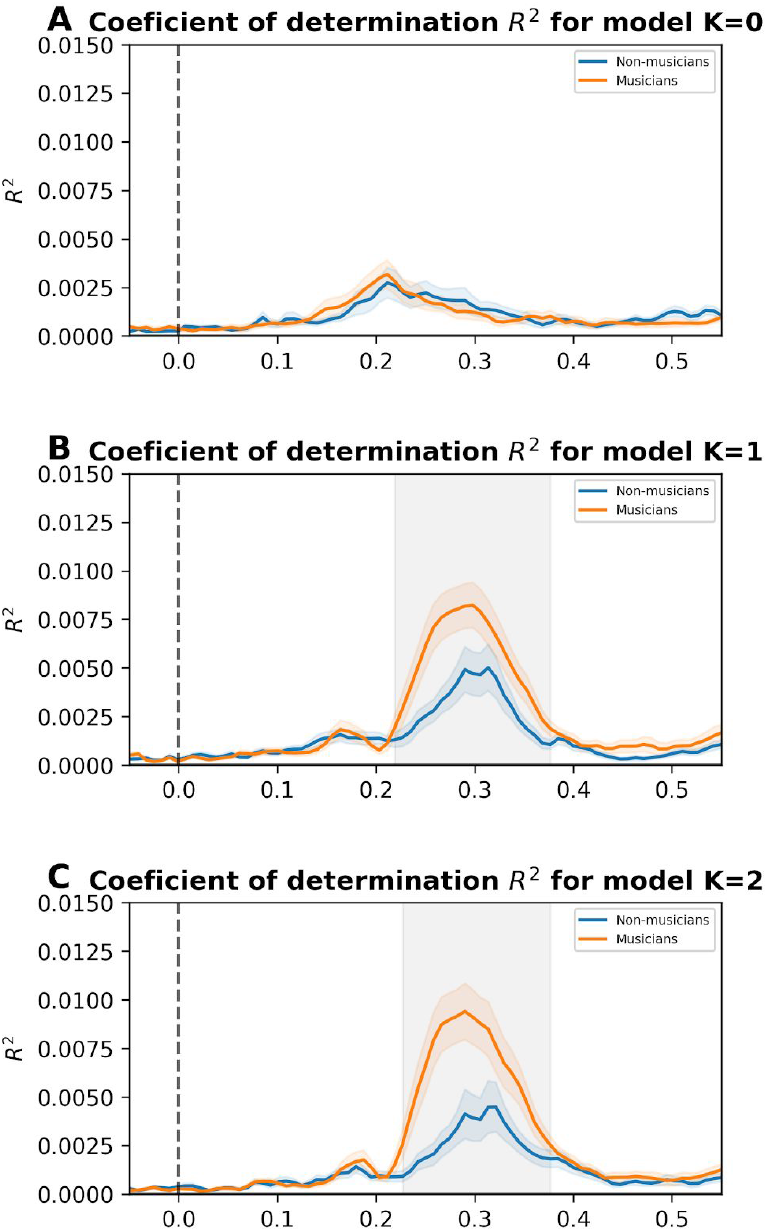
Coefficient of determination R^2^ of the linear regression between model surprise and SC amplitude for musicians and non-musicians. **A.** Coefficient of determination R^2^ of the linear regression between surprise from the simplest model (K=0) and SC amplitude for musicians and non-musicians. There is no significant difference between groups (p > 0.05, cluster-corrected) **B.** R^2^ of the linear regression between the model (K=1) surprise and SC amplitude for musicians and non-musicians. The difference between the groups is significant (p < 0.05) in a late time window, between 220 and 380 ms (shown in grey). **C.** R^2^ of the linear regression between the most complex model (K = 2) surprise and SC amplitude for musicians and non-musicians. The difference between the groups is significant (p < 0.05) in the late time window, between 220 and 380 ms (shown in grey). Colored shaded areas represent standard error of the mean (s.e.m.).

**Figure Supp. 5.**
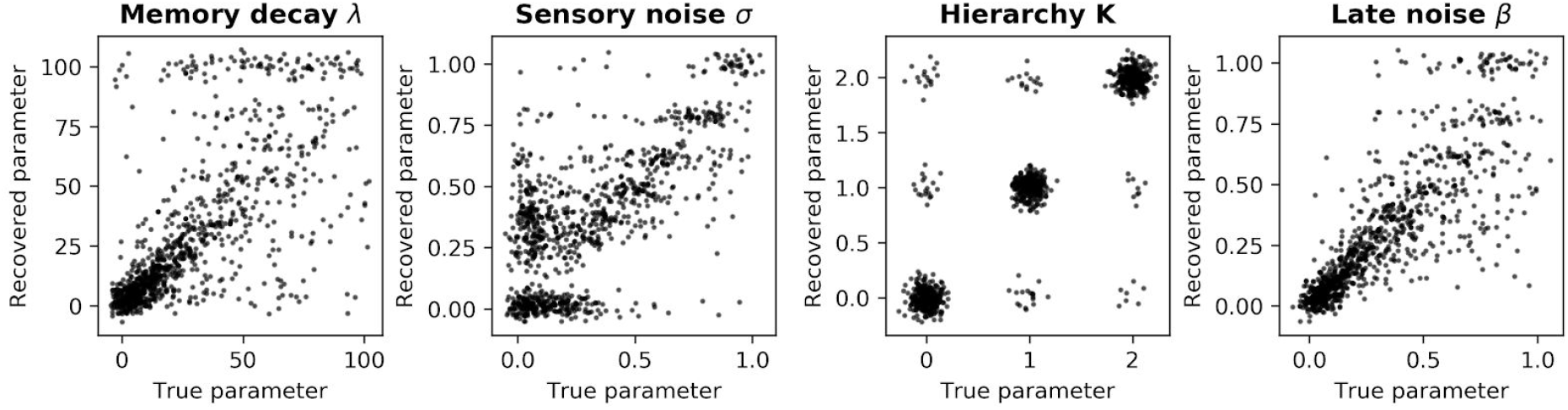
Parameter recovery analysis. We generated synthetic data for 10^3^ models with random parameters (*λ* exponential between 0 and 100, *σ*, exponential between 0 and 1, K, uniform choice between 0, 1 and 2, β exponential between 0 and 1). Recovered parameters from these synthetic data using our procedure were close to the original parameters (spearman *ϱ*_*λ*_ = 0.78, *ϱ*_*σ*_ = 0.62, *ϱ*_K_= 0.90, *ϱ*_β_ = 0.92).

## Bibliography

1. Dehaene, S., Meyniel, F., Wacongne, C., Wang, L. & Pallier, C. The neural representation of sequences: from transition probabilities to algebraic patterns and linguistic trees. Neuron 88, 2–19 (2015).

2. Summerfield, C. & de Lange, F. P. Expectation in perceptual decision making: neural and computational mechanisms. Nat. Rev. Neurosci. 15, 745–756 (2014).

3. Skinner, B. F. Science and Human Behavior. (1953).

4. Kuhl, P. K. Early language acquisition: cracking the speech code. Nat. Rev. Neurosci. 5, 831–843 (2004).

5. Romberg, A. R. & Saffran, J. R. Statistical learning and language acquisition. Wiley Interdiscip. Rev. Cogn. Sci. 1, 906–914 (2010).

6. Saffran, J. R., Aslin, R. N. & Newport, E. L. Statistical learning by 8-month-old infants. Science 274, 1926–1928 (1996).

7. Koelsch, S. Music-syntactic processing and auditory memory: similarities and differences between ERAN and MMN. Psychophysiology 46, 179–190 (2009).

8. Francois, C. & Schön, D. Musical expertise boosts implicit learning of both musical and linguistic structures. Cereb. Cortex 21, 2357–2365 (2011).

9. François, C., Tillmann, B. & Schön, D. Cognitive and methodological considerations on the effects of musical expertise on speech segmentation. Ann. N. Y. Acad. Sci. 1252, 108–115 (2012).

10. Pearce, M. T., Ruiz, M. H., Kapasi, S., Wiggins, G. A. & Bhattacharya, J. Unsupervised statistical learning underpins computational, behavioural, and neural manifestations of musical expectation. Neuroimage 50, 302–313 (2010).

11. Vuust, P., Ostergaard, L., Pallesen, K. J., Bailey, C. & Roepstorff, A. Predictive coding of music--brain responses to rhythmic incongruity. Cortex 45, 80–92 (2009).

12. Daikoku, T. Neurophysiological markers of statistical learning in music and language: hierarchy, entropy, and uncertainty. Brain Sci. 8, (2018).

13. Kolossa, A., Fingscheidt, T., Wessel, K. & Kopp, B. A model-based approach to trial-by-trial p300 amplitude fluctuations. Front. Hum. Neurosci. 6, 359 (2012).

14. Mars, R. B. et al. Trial-by-trial fluctuations in the event-related electroencephalogram reflect dynamic changes in the degree of surprise. J. Neurosci. 28, 12539–12545 (2008).

15. Meyniel, F., Maheu, M. & Dehaene, S. Human Inferences about Sequences: A Minimal Transition Probability Model. PLoS Comput. Biol. 12, e1005260 (2016).

16. Squires, K. C., Wickens, C., Squires, N. K. & Donchin, E. The effect of stimulus sequence on the waveform of the cortical event-related potential. Science 193, 1142–1146 (1976).

17. Siegelman, N., Bogaerts, L., Christiansen, M. H. & Frost, R. Towards a theory of individual differences in statistical learning. Philos. Trans. R. Soc. Lond. B. Biol. Sci 372, (2017).

18. Siegelman, N., Bogaerts, L. & Frost, R. Measuring individual differences in statistical learning: Current pitfalls and possible solutions. Behav. Res. Methods 49, 418–432 (2017).

19. Aramaki, M., Kronland-Martinet, R., Voinier, T. & Ystad, S. A percussive sound synthesizer based on physical and perceptual attributes. Computer Music Journal 30, 32–41 (2006).

20. Chase, H. W., Swainson, R., Durham, L., Benham, L. & Cools, R. Feedback-related negativity codes prediction error but not behavioral adjustment during probabilistic reversal learning. J. Cogn. Neurosci. 23, 936–946 (2011).

21. Maheu, M., Dehaene, S. & Meyniel, F. Brain signatures of a multiscale process of sequence learning in humans. elife 8, (2019).

22. Shannon, C. E. A mathematical theory of communication. Bell System Technical Journal 27, 379–423 (1948).

23. de Cheveigné, A. et al. Multiway canonical correlation analysis of brain data. Neuroimage 186, 728–740 (2019).

24. Clark, A. Whatever next? Predictive brains, situated agents, and the future of cognitive science. Behav. Brain Sci. 36, 181–204 (2013).

25. Friston, K. The free-energy principle: a unified brain theory? Nat. Rev. Neurosci. 11, 127–138 (2010).

26. Hochstein, S. & Ahissar, M. View from the top: hierarchies and reverse hierarchies in the visual system. Neuron 36, 791–804 (2002).

27. Näätänen, R. The mismatch negativity: a powerful tool for cognitive neuroscience. Ear Hear. 16, 6–18 (1995).

28. Micheyl, C., Delhommeau, K., Perrot, X. & Oxenham, A. J. Influence of musical and psychoacoustical training on pitch discrimination. Hear. Res. 219, 36–47 (2006).

29. Spiegel, M. F. & Watson, C. S. Performance on frequency-discrimination tasks by musicians and nonmusicians. J. Acoust. Soc. Am. 76, 1690–1695 (1984).

30. Kuman, P. V., Rana, B. & Krishna, R. Temporal processing in musicians and non-musicians. J Hear Sci (2014).

31. Patel, A. D. Why would Musical Training Benefit the Neural Encoding of Speech? The OPERA Hypothesis. Front. Psychol. 2, 142 (2011).

32. Paavilainen, P. The mismatch-negativity (MMN) component of the auditory event-related potential to violations of abstract regularities: a review. Int. J. Psychophysiol. 88, 109–123 (2013).

33. Herholz, S. C., Boh, B. & Pantev, C. Musical training modulates encoding of higher-order regularities in the auditory cortex. Eur. J. Neurosci. 34, 524–529 (2011).

34. Paraskevopoulos, E., Kuchenbuch, A., Herholz, S. C. & Pantev, C. Musical expertise induces audiovisual integration of abstract congruency rules. J. Neurosci. 32, 18196–18203 (2012).

35. Tervaniemi, M., Rytkönen, M., Schröger, E., Ilmoniemi, R. J. & Näätänen, R. Superior formation of cortical memory traces for melodic patterns in musicians. Learn. Mem. 8, 295–300 (2001).

36. Fujioka, T., Trainor, L. J., Ross, B., Kakigi, R. & Pantev, C. Musical training enhances automatic encoding of melodic contour and interval structure. J. Cogn. Neurosci. 16, 1010–1021 (2004).

37. Koelsch, S., Schmidt, B.-H. & Kansok, J. Effects of musical expertise on the early right anterior negativity: an event-related brain potential study. Psychophysiology 39, 657–663 (2002).

38. Roads, C. & Strawn, J. The computer music tutorial. (1996).

39. Girshick, A. R., Landy, M. S. & Simoncelli, E. P. Cardinal rules: visual orientation perception reflects knowledge of environmental statistics. Nat. Neurosci. 14, 926–932 (2011).

40. Ernst, M. O. & Banks, M. S. Humans integrate visual and haptic information in a statistically optimal fashion. Nature 415, 429–433 (2002).

41. Drugowitsch, J., Wyart, V., Devauchelle, A.-D. & Koechlin, E. Computational precision of mental inference as critical source of human choice suboptimality. Neuron 92, 1398–1411 (2016).

42. Kahneman, D. & Tversky, A. Prospect theory. an analysis of decision making under risk. (US Dept of the Navy, 1977). doi:10.21236/ADA045771

43. Brunton, B. W., Botvinick, M. M. & Brody, C. D. Rats and humans can optimally accumulate evidence for decision-making. Science 340, 95–98 (2013).

44. Kaufman, M. T. & Churchland, A. K. Cognitive neuroscience: sensory noise drives bad decisions. Nature 496, 172–173 (2013).

45. Osborne, L. C., Lisberger, S. G. & Bialek, W. A sensory source for motor variation. Nature 437, 412–416 (2005).

46. Sutton, R. S. & Barto, A. G. Reinforcement Learning: An Introduction. IEEE Trans. Neural Netw. 9, 1054–1054 (1998).

47. Acerbi, L., Vijayakumar, S. & Wolpert, D. M. On the origins of suboptimality in human probabilistic inference. PLoS Comput. Biol. 10, e1003661 (2014).

48. Dayan, P. Rationalizable irrationalities of choice. Top. Cogn. Sci. 6, 204–228 (2014).

49. Beck, J. M., Ma, W. J., Pitkow, X., Latham, P. E. & Pouget, A. Not noisy, just wrong: the role of suboptimal inference in behavioral variability. Neuron 74, 30–39 (2012).

50. Renart, A. & Machens, C. K. Variability in neural activity and behavior. Curr. Opin. Neurobiol. 25, 211–220 (2014).

51. Koelsch, S., Vuust, P. & Friston, K. Predictive processes and the peculiar case of music. Trends Cogn Sci (Regul Ed) 23, 63–77 (2019).

52. Perruchet, P. & Pacton, S. Implicit learning and statistical learning: one phenomenon, two approaches. Trends Cogn Sci (Regul Ed) 10, 233–238 (2006).

53. Brady, T. F., Konkle, T. & Alvarez, G. A. Compression in visual working memory: using statistical regularities to form more efficient memory representations. J. Exp. Psychol. Gen. 138, 487–502 (2009).

54. Ahissar, M. & Hochstein, S. Task difficulty and the specificity of perceptual learning. Nature 387, 401–406 (1997).

55. Schellenberg, E. G. Correlation = causation? Music training, psychology, and neuroscience. Psychol. Aesthet. Creat. Arts (2019). doi:10.1037/aca0000263

56. François, C., Chobert, J., Besson, M. & Schön, D. Music training for the development of speech segmentation. Cereb. Cortex 23, 2038–2043 (2013).

57. Flaugnacco, E. et al. Music training increases phonological awareness and reading skills in developmental dyslexia: A randomized control trial. PLoS ONE 10, e0138715 (2015).

58. Hidalgo, C., Falk, S. & Schön, D. Speak on time! Effects of a musical rhythmic training on children with hearing loss. Hear. Res. 351, 11–18 (2017).

59. Hidalgo, C., Pesnot-Lerousseau, J., Marquis, P., Roman, S. & Schön, D. Rhythmic training improves temporal anticipation and adaptation abilities in children with hearing loss during verbal interaction. J. Speech Lang. Hear. Res. 62, 3234–3247 (2019).

60. Sihvonen, A. J. et al. Music-based interventions in neurological rehabilitation. Lancet Neurol. 16, 648–660 (2017).

61. Bates, D., Mächler, M., Bolker, B. & Walker, S. Fitting linear mixed-effects models using lme4. arXiv preprint arXiv:1406.5823 (2014).

62. Pearce, M. T. & Wiggins, G. A. Auditory expectation: the information dynamics of music perception and cognition. Top. Cogn. Sci. 4, 625–652 (2012).

63. Gramfort, A. et al. MEG and EEG data analysis with MNE-Python. Front. Neurosci. 7, 267 (2013).

64. Brooks, S., Gelman, A., Carlin, J. B., Stern, H. S. & Rubin, D. B. Bayesian Data Analysis. The Statistician 45, 266 (1996).

